# The differing prevalences of propionate and butyrate-producing bacteria in the human gut microbiota

**DOI:** 10.1101/2023.11.27.568948

**Authors:** Rebecca Christensen, Yu Han Daisy Wang, Markus Arnoldini, Jonas Cremer

## Abstract

Propionate and butyrate are major fermentation products released during the anaerobic growth of bacteria in the large intestine with strong implications on host health. While the different metabolic pathways leading to the release of these products are biochemically well characterized, less is known about their relative abundances across hosts and conditions. Here, we introduce a bioinformatics pipeline which connects pathway analyses, gene identification via sequence homology, and the screening of metagenomic samples to systematically identify the abundances of propionate producing pathways across individuals and in relation to butyrate producing pathways. We found that on average 36% of all genomes of a gut microbiota carried propionate producing pathways with the sodium-pumping succinate pathway being the most prevalent. This pathway abundance was anti-correlated to the abundance of butyrate pathways and greatly depended on host physiology. For example, propionate pathway abundance varied strongly among infants with a substantially higher abundance in vaginally than C-section born infants, increased in early adulthood, and decreased again with higher ages. This is in strong contrast to the known variation of butyrate pathway abundance, which is close to zero in infants and steadily increases with age. These results highlight the contrasting prevalence of both fermentation products at the individual level with shifts in microbiota composition resulting in strong consequences on the amount of butyrate and propionate available to the host.

## Introduction

Interactions between the human gut microbiota and its host are driven by the exchange of different molecules. In the anoxic environment of the large intestine, the microbiota is dominated by bacteria which catabolize dietary components, mostly complex carbohydrates, via anaerobic fermentation. To supply energy while maintaining an oxidation-reduction balance, microbial fermentation leads to the secretion of large amounts of acetate, butyrate, propionate and other molecules. Importantly, gut bacteria vary strongly in the excretion of these different fermentation products. While acetate is ubiquitously produced by most microbial fermenters in the human gut, butyrate as well as propionate are only produced by a fraction of gut microbes (Louis & Flint, 2017). Furthermore, while propionate and butyrate are both absorbed by the gut epithelium, they are differently processed and utilized by the host. Particularly, butyrate is largely consumed in the colonic epithelium where it serves as a major energy source for colonocytes (Cummings et al., 1987; Donohoe et al., 2011; McNeil et al., 1978). In contrast, propionate moves through the circulatory system and is thought to be primarily utilized in the liver to support cellular respiration (Cummings et al., 1987; Donohoe et al., 2011; Weitkunat et al., 2016). Propionate has also been implicated as an ameliorative force in obesity and was suggested to reduce weight gain and promote the release of anorectic gut hormones in colonocytes (Chambers et al., 2015). Taken together, these variations in bacterial fermentation combined with the differential utilization of each fermentation product by the host suggests a specific mechanism by which the composition of the microbiota strongly affects host health.

In line with these ideas, several studies have reported a change in host health coupled with fermentation product secretion and microbiota composition (Machiels et al., 2014; Mirzaei et al., 2021; Wang et al., 2012; Wu et al., 2020). Both butyrate and propionate have been indicated to modulate inflammatory as well as immune processes and promote antitumor effects (Miller et al., 2005; Mirzaei et al., 2021; Nastasi et al., 2015; Tang et al., 2011; Zapolska-Downar et al., 2004; Zapolska-Downar & Naruszewicz, 2009). Additionally, the prevalence of common butyrate producing species was found to be diminished in individuals with colorectal cancer, ulcerative colitis, and type 2 diabetes mellitus (Machiels et al., 2014; Wang et al., 2012; Wu et al., 2020). In conclusion, elucidating the varying capability of the microbiota composition to produce butyrate and propionate is therefore a critical step towards a broader understanding of the relationship between the gut microbiota and human health.

Previous studies have attempted to directly measure the prevalence of propionate and butyrate in the human gut, though this experimental approach represents a major challenge. For example, the rapid uptake of short chain fatty acids within the large intestine renders fecal samples, the most common sample type for the gut microbiota, unsuitable. Furthermore, concentrations can vary strongly in space and time with a strong dependence on fluid turnover and food intake by the host, two quantities which vary highly throughout the day. Thus, even local concentration measurements in the proximal large intestine provide limited insights into the amount of fermentation products released by bacteria. As an alternative, the variation in fermentation products released with microbiota composition can be estimated by the taxonomic identification of experimentally established butyrate and propionate producing species in microbiota samples. Coupled with the quantitative measurement of fermentative rates, this approach can illustrate the variation of fermentation product release with microbiota composition, an approach we introduce in a different study (Arnoldini et al., in preparation). However, this approach relies on a sufficiently representative majority of bacterial species in a microbiota sample to be experimentally characterized. It thus has limitations when exploring the capabilities of butyrate and propionate productions across very diverse microbiota samples and is best complemented by a bioinformatics approach which is independent of experimental characterizations and can, in parallel, analyze the metabolic capabilities of hundreds or even thousands of microbiota samples. This third type of approach particularly identifies the abundance of different fermentation producing pathways in metagenomic data as proxy for the capability of the microbiota to release different fermentation products. Importantly, the functions of these pathway genes are often ambiguous and utilized in different pathways. Thus, the consideration of a single metabolic gene alone can be misleading when estimating the capability of the microbiota to release fermentation products. Instead, the quantification of entire pathway abundances in metagenomic samples can offer a more accurate determination of fermentation capabilities across different microbiota samples. Vital et al. demonstrated this pathway abundance approach by characterizing and quantifying the abundances of different butyrate producing pathways in the human gut (Vital et al., 2014, 2017). The results emphasize a strong variation of butyrate pathway abundances across different microbiome samples. While based on only single pathway genes, a recent bioinformatics study has furthermore demonstrated a similarly strong variation in the abundance of propionate producing pathways (Medina et al., 2021). From these studies it is clear that there exists a wide variation in butyrate and propionate pathway abundance. However, we currently lack a systematic pathway analysis for propionate, and the relative capability of the microbiota to produce both of these major metabolites remains poorly investigated. Notably, as gut species commonly harbor only one of eight butyrate or propionate producing pathways, we expect strong anticorrelations in the capabilities of a microbiota to produce either propionate or butyrate. In this work, we thus quantify the relative abundances of propionate and butyrate pathways by developing a scalable pipeline that maps metagenomic samples of the human gut to our curated catalog of pathway-specific genes involved in propionate and butyrate production. We apply this pipeline to analyze the variation of propionate and butyrate pathway abundance across a variety of hosts, specifically discussing the roles of host age and inflammatory bowel disease.

## Results

### Pathway abundance analysis

Four fermentative pathways lead to the microbial production of butyrate and propionate, respectively (**Fig. 1A**) (Gonzalez-Garcia et al., 2017; Louis & Flint, 2017). Here we name them after the metabolite entry point of each pathway. Pathways for butyrate production include the glutarate (Glu), the acetyl-CoA (Ace), the lysine (Lys), and the 4-aminobutyrate/succinate pathway (4-Ami). Pathways for propionate production include the propanediol (Pro), the acrylate (Acr), and the succinate pathway. The latter is further subdivided into the Wood-Werkman cycle (WWC) and the sodium-pumping pathway (SP). To estimate the relative abundances of each pathway from metagenomics data, we developed a bioinformatics pipeline which, similar to previous approaches (Vital et al., 2014, 2017), combines the construction of profile Hidden Markov models from well characterized model strains (**Supplementary Table S1**) to identify pathway genes, the compilation of a gut-specific catalog of pathway genes using a genome collection covering over 1,000 gut bacteria (Forster et al., 2019), and the screening of metagenomic samples for these genes in different metagenomic samples from the microbiota of different individuals. The major steps of the pipeline are outlined in **Fig. 1B** with further rationalization and implementation details provided in the **Materials and Methods** section and on the GitHub repository. To confirm the validity of this pipeline we first tested the ability of the pipeline to accurately predict the pathway presence in experimentally well characterized model species. We found good agreement with the previous reports. Particularly, Clostridia species known for butyrate production were found to maintain the butyrate pathway genes while propionate pathways were detected in many Bacteroides strains. All identified propionate and butyrate producers are listed in **Supplementary Table S2**. To estimate pathway abundance in microbiota samples, we then ran the pipeline to obtain the percentage of genomes that carry the eight propionate and butyrate producing pathways from metagenomic data.

**Figure 1.**
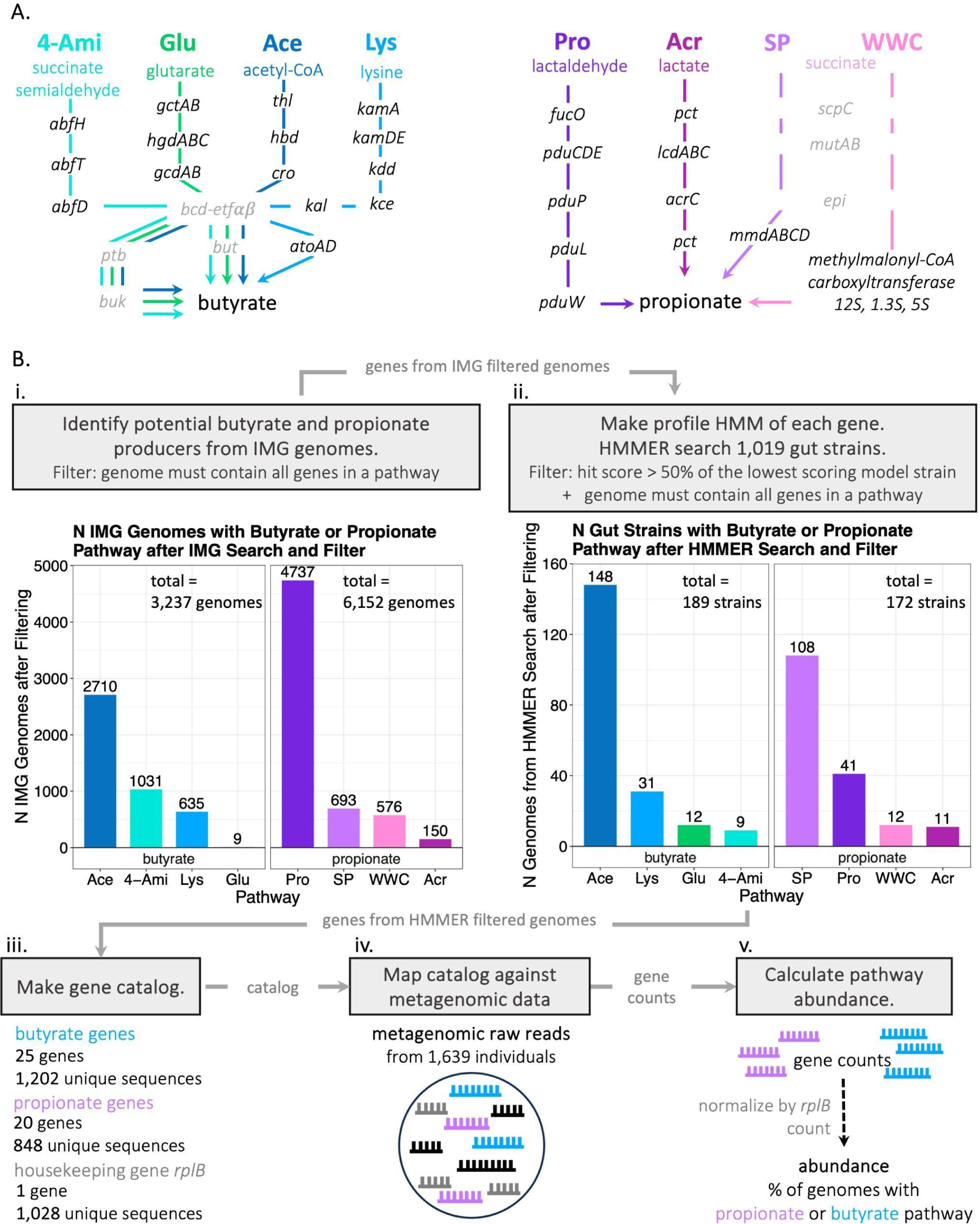
Quantification of butyrate and propionate pathway abundances. **A)** Different butyrate and propionate producing metabolic pathways, named according to their metabolite entry point (colored labels, name abbreviations in bold). Catalyzing genes are shown in black/gray. Genes in gray are utilized by multiple pathways. **B)** Overview of bioinformatics pipeline to quantify pathway abundances. Identification of potential butyrate and propionate producing strains from the Integrated Microbial Genomes (IMG) database **(i)**, which were used to build profile Hidden Markov models to identify homologous sequences (HMMER) in genomes of gut bacteria **(ii)**. Bar charts show the occurrence of each pathway in all genomes from IMG (left) and genomes from gut bacteria (right). Construction of gene catalog **(iii)** to enumerate abundances of pathway genes in metagenomic samples **(iv)**. Normalization of the number of pathway genes found in a metagenomic sample against the housekeeping gene *rplB* to calculate the relative abundance of genomes containing each pathway **(v)**.

### Variation of pathway abundance in healthy humans

To analyze the variation in propionate and butyrate pathway abundance across individuals, we first analyzed available metagenomic data from over 1,500 healthy individuals from a variety of geographical regions and ages (**SI Text S1**) (Asnicar et al., 2017, 2021; Bäckhed et al., 2015; Lloyd-Price et al., 2019; Schirmer et al., 2018).

Overall, butyrate and propionate pathways were highly abundant, with on average 64% of all genomes carrying one of the pathways. Present in 38% of genomes in the microbiota, propionate pathways were on average more abundant than butyrate producing ones with the abundance being roughly normally distributed around the average. Very few individuals contained little to no propionate pathways (i.e. we detected a propionate pathway abundance of less than 1% of genomes in less than 1% of metagenomic samples) (**Fig. 2A**, pink distribution). The total abundance of all propionate pathways was overwhelmingly dominated by the SP pathway, followed by the Pro pathway. The remaining two pathways were rarely observed (**Fig. 2B**).

**Figure 2.**
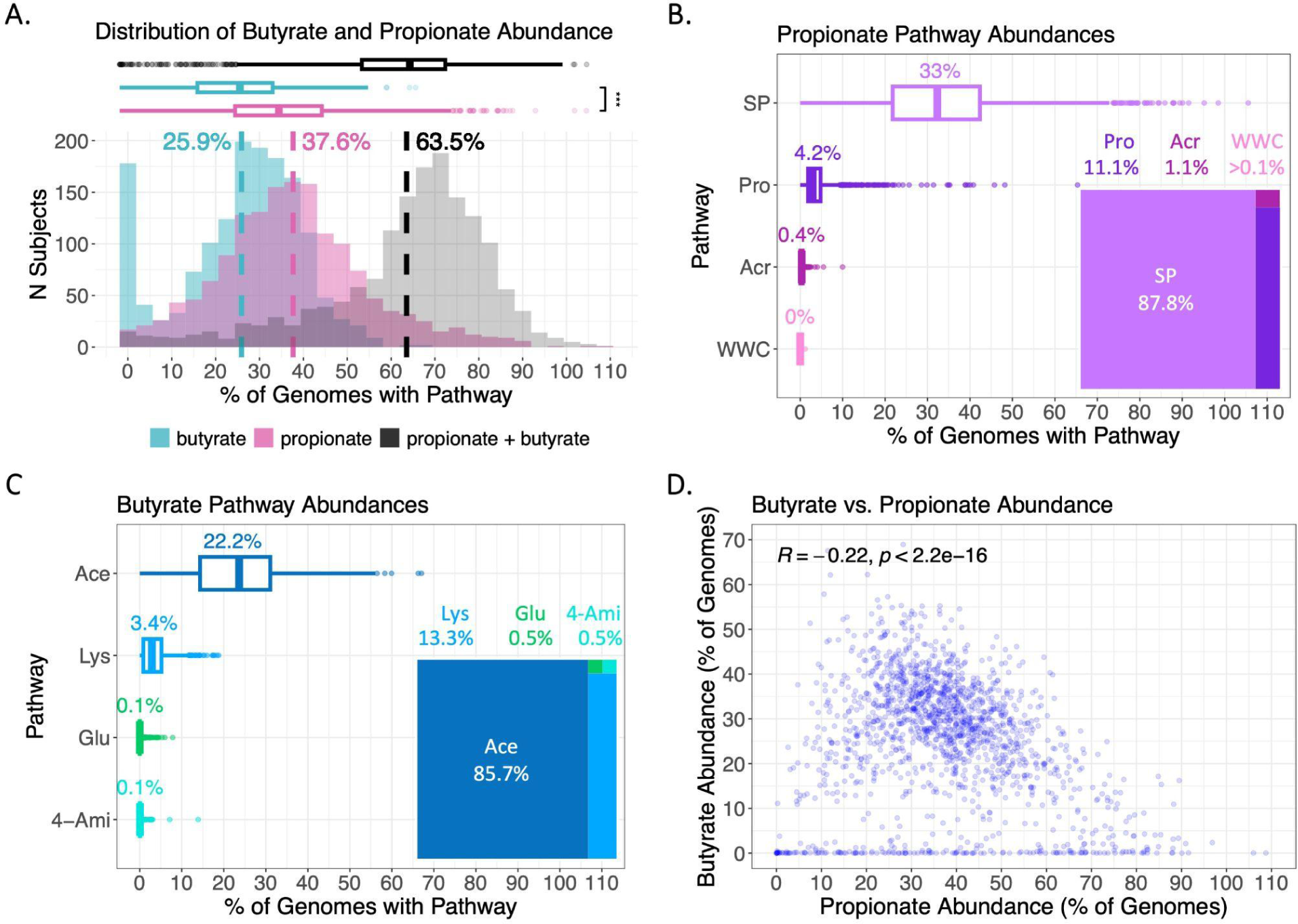
Variation of propionate and butyrate pathway abundance in healthy individuals. **A)** Distribution of the percentage of genomes containing butyrate pathways, propionate pathways, and the sum. The average fraction of genomes containing propionate pathways is significantly higher than that of butyrate. Dashed lines indicate means. **B and C)** Abundance of the different propionate and butyrate pathways. Numbers above box plots indicate means. Insets illustrate the relative abundances of different butyrate/propionate pathways in comparison to the total butyrate/propionate pathway abundance. Data from 1,626 metagenomic samples from *n* = 1,535 healthy individuals (Asnicar et al., 2017, 2021; Bäckhed et al., 2015; Lloyd-Price et al., 2019; Schirmer et al., 2018). Significance in A-C,confirmed by Wilcoxon rank sum test, *p*-value < 0.05. **D)** Relation between propionate and butyrate abundance. Spearman’s rank correlation coefficient shown.

Butyrate pathway abundance also varied strongly across individuals, in agreement with previous studies (Vital et al., 2014, 2017) (**Fig. 2A**, teal distribution). Abundance was typically lower than that of propionate with a 12% reduction in the average percentage of genomes containing butyrate pathways. Furthermore, in contrast to propionate, many individuals harbored microbiomes containing little to no butyrate pathway genes (i.e. in 10% of individuals we detected a butyrate pathway abundance of less than 1%). The Ace pathway overwhelmingly dominated butyrate pathway abundance (Vital et al., 2014, 2017) with the Lys pathway being the second most abundant (**Fig. 2C**).

The abundances of butyrate and propionate pathways were furthermore anticorrelated (**Fig. 2D**). Particularly, butyrate pathway abundance was consistently relatively low in samples where propionate pathway abundance was relatively high. Given that gut bacteria commonly harbor only one of the eight propionate and butyrate pathways (**SI Text S1**), this anticorrelation is expected if the overall abundance of propionate and butyrate pathways is high. This further underlines the importance of butyrate and propionate production for gut bacteria to maintain a physiological redox balance and enable anaerobic growth.

In summary, the pipeline reveals differing variations in butyrate and propionate pathway abundance across microbiota samples from different individuals, indicating a strong variation in the amount of different fermentation products released and highlighting the need for further studies to better understand this variation.

### Variation with age

To further put these variations in pathway abundance into perspective, we next analyzed the role of age. Previous studies have shown age to be a driving factor to changes in gut microbiome composition (Asnicar et al., 2017; Ghosh et al., 2022). To analyze the variation of pathway abundance with age, we categorized subjects into six age groups which were shown to be biologically relevant to distinct gut microbiota composition (Ghosh et al., 2022).

While butyrate pathways were hardly detected in infants, the abundance of butyrate pathways drastically increased after six months, in line with Vital et al (Vital et al., 2014, 2017). Abundance consistently increased with age, reaching a peak value (an average butyrate pathway abundance of 31.2% of genomes) in adults over 20 years old (**Fig. 3A**). Furthermore, while the dominant Ace pathway was already abundant in children 6 months to 3 years old, the Lys pathway abundance only increased in individuals from 3 to 20 years old (**Supplementary Fig. S1A**).

**Figure 3.**
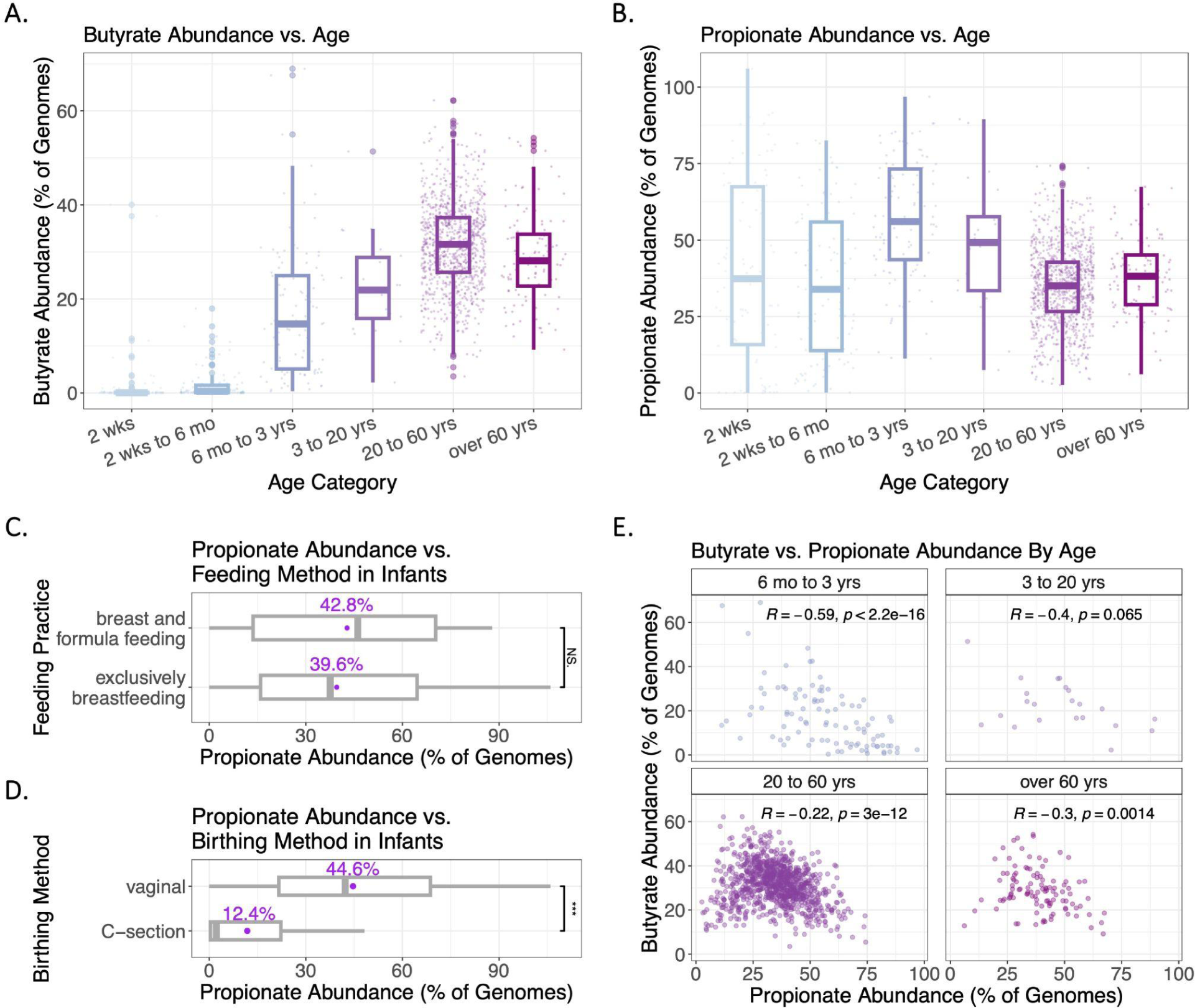
Butyrate and propionate abundance variation by age. **A and B)** Butyrate and propionate pathway abundance by age category. **C and D)** Propionate pathway abundance by feeding and birthing methods for infants less than 2 weeks old. Numbers above box plots indicate mean values. **E)** Variation in the strength of the anticorrelation between propionate and butyrate abundance in different age groups. Spearman’s rank correlation coefficient shown. Data from 1,513 metagenomic samples from *N* = 1,415 healthy individuals with available data on age (Asnicar et al., 2017, 2021; Bäckhed et al., 2015; Lloyd-Price et al., 2019; Schirmer et al., 2018). Significance in A, B, and D confirmed by Wilcoxon rank sum test, *p*-value < 0.05.

These results are in strong contrast to propionate pathway abundance. Infants already displayed a high abundance, albeit with large variation (**Fig. 3B**). While this variation in propionate pathway abundance decreased with increasing age, the average abundance increased, peaking at an average of 56.7% of genomes in children between 6 months and 3 years of age, before dropping again in adults. Throughout all age groups, SP and Pro remained the most prevalent propionate pathways (**Supplementary Fig. S1B**).

Given the particularly strong variation of propionate pathway abundances in infants, we investigated abundance in relation to different feeding practices and birthing methods for infants under 2 weeks old. While we did not detect a difference in abundance between exclusively breastfed versus breast plus formula fed infants (**Fig. 3C**), there was a significant difference in abundance between birthing methods (**Fig. 3D**). Infants born vaginally displayed an average propionate pathway abundance of 44.6% of genomes, while those born via Cesarean section had a substantially lower abundance of 12.4% of genomes on average.

We further analyzed the anticorrelation between propionate and butyrate pathway abundance in different age groups. While the strength of this anticorrelation varied between different age groups and was strongest in children and adolescents aged 6 months to 20 years, (**Fig. 3E**) an anticorrelation was observed in all age groups 6 months or older. Overall, these findings emphasize further changes in the gut microbiota with age characterized by a drastic increase in butyrate production after six months in line with a shift in diet. Given that bacterial genomes carry either butyrate or propionate pathways, the consistent anticorrelation across all age groups 6 months or older also indicates again the overall importance of propionate and butyrate production in maintaining a cellular redox balance and enabling fermentative growth.

### Variation with intestinal health

Next, we analyzed the correlation between gut health and pathway abundance in a cohort of both healthy individuals and individuals with Inflammatory Bowel Disease (IBD), specifically persons diagnosed with either Crohn’s Disease (CD) or Ulcerative Colitis (UC). We found significant differences in propionate and butyrate pathway abundances in relation to IBD status. First, butyrate pathway abundance was significantly higher in healthy individuals compared to those with IBD, in line with previous observations (Vital et al., 2017) (one-sided Wilcoxon rank sum, *p*-value = 2. 21 · 10^−7^). Second, propionate pathway abundance showed the opposite trend and was significantly higher in individuals with IBD (one-sided Wilcoxon rank sum, *p*-value = 2. 86 · 10^-16^) (**Fig. 4A**). Similar to the overall pathway variation (**Fig. 1** above), propionate and butyrate pathway abundances were relatively similar in healthy subjects with only an 11.7% difference in the average percentage of genomes. Conversely, the average propionate pathway abundance in individuals with IBD was three times higher than butyrate pathway abundance. These trends were mostly driven by changes in the phylum-level abundance of Bacteroidota and Bacillota, while *Escherichiia coli* and other propionate producing members of Pseudomonadota showed no significant changes in abundance (**Fig. 4BC**).

**Figure 4.**
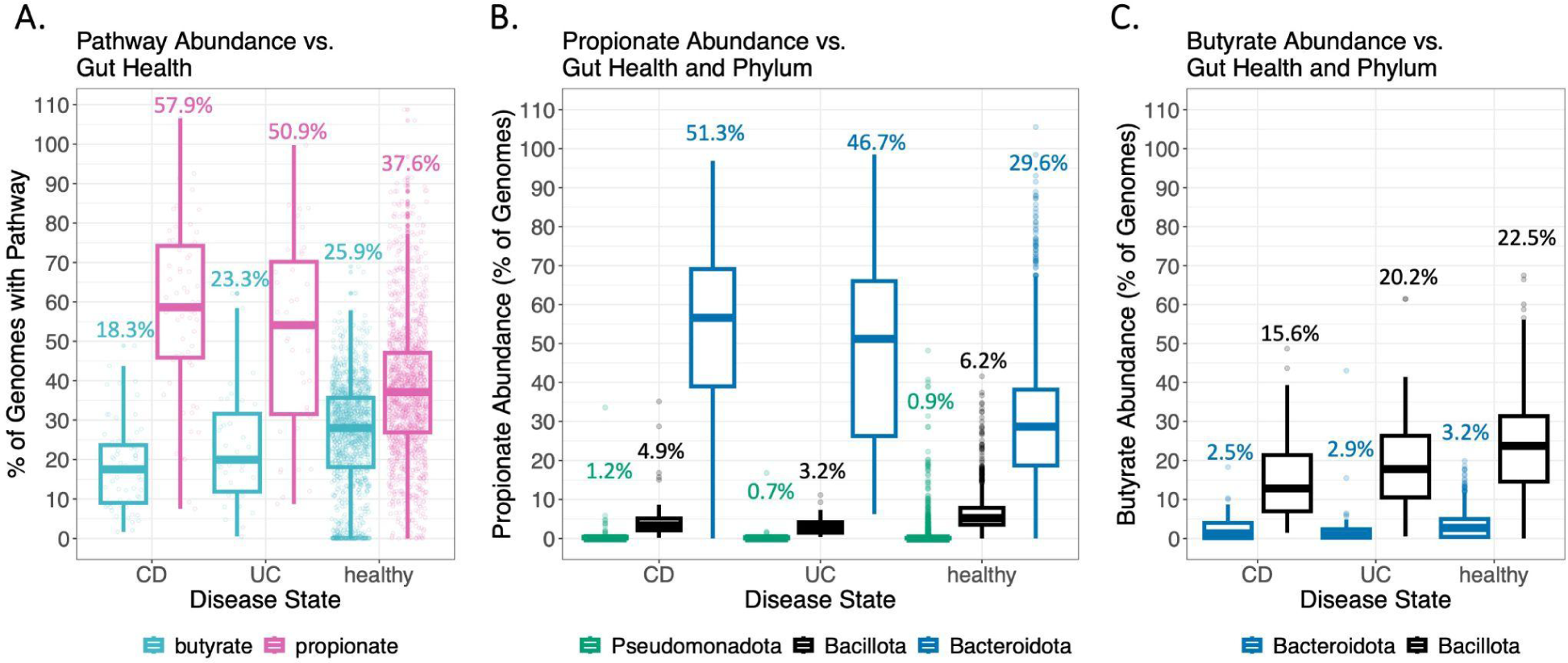
Variation in propionate and butyrate abundance in healthy versus IBD-diagnosed individuals. **A)** Opposite trends in the variation of butyrate and propionate abundances in healthy subjects versus individuals with Crohn’s disease or Ulcerative Colitis. **B and C)** Variation in propionate and butyrate abundance by most prevalent phyla. Numbers above box plots show means. Data from 1,973 samples from *N* = 1,639 subjects with available data on IBD diagnosis (*n* = 1,535 healthy subjects and *n =* 104 subjects with IBD, including *n =* 65 subjects with CD and *n =* 39 subjects with UC) (Asnicar et al., 2017, 2021; Bäckhed et al., 2015; Lloyd-Price et al., 2019; Schirmer et al., 2018). Significance between IBD and healthy groups for both propionate and butyrate pathway abundances in A confirmed by one-sided Wilcoxon rank sum test, *p-*value < 0.05.

## Discussion

The massive amount of fermentation products exchanged between the gut microbiota and the host constitutes an important mechanistic avenue in shaping microbe-host interactions. As different gut bacteria release different fermentation products with differing effects on host physiology, the effect of microbial fermentation on host health may be highly dependent on the composition of the gut microbiota. Towards exploring this link, we developed a bioinformatics pipeline that estimates in metagenomic samples from different individuals the relative abundances of butyrate and propionate producing fermentation pathways. We accomplished this by identifying pathway genes from potential propionate and butyrate producers, generating profile Hidden Markov models to detect all pathway genes, mining metagenomic data from human microbiota samples, and calculating the percentage of genomes containing each pathway within a sample as a proxy for pathway abundance. We particularly applied this scalable bioinformatics pipeline to study the variation of pathway abundances across different age groups and individuals with gut inflammation.

On average, 63.5% of genomes carried one of the eight butyrate and propionate producing pathways, highlighting the importance of these pathways for fermentative growth in the gut. The abundance of propionate producing pathways was greater than that of butyrate, albeit with strong variation. Notably, our approach also allowed us to relate propionate and butyrate pathway abundances, going beyond previous studies that analyzed propionate and butyrate production independently (Medina et al., 2021; Reichardt et al., 2014; Vital et al., 2014, 2017). Our analysis showed a contrary relation between propionate pathway abundance and butyrate pathway abundance in different samples. Particularly, when the overall abundance of propionate was high, the butyrate pathway abundances was consistently low. This suggests a competition between butyrate- and propionate-producing microbes as expected from the overlap in resource consumption, specifically for carbohydrate and amino acid fermentation.

Pathway abundances also varied greatly with age. Particularly, while butyrate pathway abundance was insubstantial in infants, it steadily increased with age. In contrast, propionate abundance was higher among infants but with strong fluctuations which decreased with age. Further investigation revealed a strong correlation between propionate pathway abundance and birthing method in infants, as vaginally born infants had significantly higher abundances of propionate pathways compared to those born via C-section.

We further found that butyrate pathway abundance was markedly higher in healthy individuals compared to individuals with IBD in agreement with previous studies (Machiels et al., 2014; Vital et al., 2017). Conversely, we obtained the opposite results for propionate, with healthy individuals displaying a lower propionate pathway abundance than individuals diagnosed with CD or UC. While no evidence has suggested that propionate producers in the human gut pose a direct detriment to gut health, the decrease in butyrate producers in individuals with IBD may open the niche in the same gut environment for propionate producers to flourish, and the establishment of a dominant propionate producing population may limit the growth of butyrate producers essential to gut health. Further analyses of the relation between propionate and butyrate pathway abundances in relation to disease progression is needed to explore this possibility and the relationship between the native microbiome with IBD etiology and onset.

Building on the analysis pipeline, other relations between host physiology, behavior, and microbiota fermentation can be studied next to better understand the role of fermentation products in shaping microbiota-host interactions. For example, we believe it would be particularly illuminating to apply this pipeline to analyze the relationship between butyrate and propionate pathway abundances to host diet. Previous research has established diet as a major determinant of microbiota composition and further linked variation in diet to drastic changes in gut microbiome composition and the onset of gastrointestinal inflammatory conditions (Beam et al., 2021; Gill et al., 2022; Malesza et al., 2021). Finally, we also hope that the bioinformatics pipeline introduced here and the focused analysis of relative pathway abundances may be further applied beyond the dissection of propionate and butyrate production. This includes the fermentative production of branched chain fatty acids utilized for the catabolism of peptides, a process which also leads to toxin release and has been suggested to be highly health detrimental in excess (Duncan et al., 2021; Macfarlane & Allison, 1986).

## Methods and Materials

### Bioinformatics pipeline to estimate relative pathway abundance in metagenomic datasets

As outlined in **Fig. 1**, our pipeline consists of five major steps which we introduce here in more detail.

Step 1: To start, we identified bacterial genomes from the Integrated Microbial Genomes database (Chen et al., 2023) that contained one of the eight fermentation pathways: First, we searched the IMG database for gene entries using entries of the Kyoto Encyclopedia of Genomes and Genes Orthology Database (Kanehisa & Goto, 2000) for each pathway gene. Based on the presence of IMG gene entries for metabolically well-characterized model strains listed in **Supplementary Table 1**, we then derived a list of required genes for each pathway: aside from *etfAB*, *acrC*, and *mmdD*, which were excluded due to lack of IMG gene entries in model strains, all genes were required to be present. Entries for *bcd*, *but*, *ptb*, and *hbd* were filtered for keywords in the description to exclude non-specific entries. We then used this list of required genes to filter 22,269 bacterial genomes available at IMG. We identified 6,152 and 3,237 bacterial genomes containing the butyrate and propionate pathways respectively, with strong variation among the prevalence of each specific pathway (**Fig. 1B (i)**).

Step 2: Next, we constructed a profile Hidden Markov Model (HMM) (Eddy, 1998) of each pathway gene using IMG genomes that contained the respective pathway. These profile HMMs were then used to HMMER search over 1,000 genomes from gut bacteria, including common gut strains (Forster et al., 2019) and model propionate and butyrate producers, to identify strains containing butyrogenic or propiogenic pathways. Hit sequences from queried strains returned by the HMMER search were filtered with a hit score cutoff of 50% of the lowest scoring model strain in accordance with an obvious score drop off. HMMER scores for each pathway were compared between strains that tested positive for multiple pathways in order to recategorize these strains as containing only one pathway. Following this analysis, only 11 strains remained categorized as containing both the Lys and Ace butyrogenic pathways. Strains that tested positive for multiple pathways were further investigated to determine the likelier pathway (**Supplementary Information - Text 1**). Of the 1,109 gut strains that were queried in the HMMER search and subsequently filtered by the hit score cutoff and the presence of all required genes, 172 strains contained a propiogenic pathway, and 189 contained a butyrogenic pathway. The most common pathways were SP for propionate, and Ace for butyrate (**Fig. 1B(ii)**).

Steps 3: From the genomes that were positive for a butyrate or propionate pathway, we compiled a gene catalog containing over 2,000 unique sequences (**Fig. 1B(iii)**).

Step 4: We then read-mapped the gene catalog to mine different metagenomic datasets for propionate and butyrate pathway genes using BOWTIE2 (Langmead & Salzberg, 2012) (**Fig. 1B(iv)**). From the obtained number of hits for each gene of a pathway gene we then determined the number of hits a pathway by accounting for the length of each pathway gene and the number genes in a pathway. We did this for each of the eight pathways. Further calculation details are provided below (**Methods and Materials - Pathway abundance count and normalization**).

Step 5: Finally, to estimate the percentage of genomes present in a metagenomic sample that contains each pathway, we normalized pathway gene counts by counts for the housekeeping gene *rplB*. This gene encodes for the rRNA binding component of the 50S ribosomal subunit and is consistently present with a copy number of 1.03 in different genomes from gut bacteria (Vital et al., 2017). Similar to the pathway counts, we determined *rplB* counts by comparing metagenomic sequences against a gene catalog of *rplB* genes derived from a HMMER search against gut bacteria with a reporting score of at least 400.

### Pathway abundance count and normalization

For each metagenomic sample the percentage of genomes that contained a specific pathway was calculated as *P_genomes_* = ∑((*h*_gene_ / *l_gene_*) / (*l_pathway_*)) / ∑(*h_housekeeping_* / *l_housekeeping_*) ⋅ 100. Here, *h*_gene_ denotes the number of obtained hits for a pathway gene, *l_gene_* is the length of that gene in nucleotides, and *l_pathway_* is the number of genes in the pathway. *h_housekeeping_* denotes the number of hits for each *rplB* sequence and *l_housekeeping_* is the *rplB* sequence length in nucleotides. For multi-pathway genes, *h*_gene_ was adjusted to reflect the proportion of single-pathway genes for each pathway in the same metagenomic sample under the assumption that the division of *h*_gene_ for multi-pathway genes is proportional to the pathway abundance, approximated by *h*_gene_ for single-pathway genes, calculated as follows: *h_gene_* = *h_o_* ⋅ (∑*h_s_* / ∑*h_m_*), with *h_o_* as the observed gene count of the multi-pathway gene, *h_s_* as the gene count for each single-pathway gene in the pathway of interest, and *h_m_* as the gene count for each single-pathway gene in all pathways. All boxplots shown throughout the manuscript are Tukey boxplots (McGill et al., 1978).

### Data Availability

The pipeline for running the HMMER search, filtering the HMMER, constructing the gene catalog, performing the BOWTIE2 read-mapping, and calculating pathway abundances is available at https://github.com/rchristensen26/Butyrate_Propionate_Comparative_Analysis. Included in the pipelines are all input files, accession numbers for metagenomic data analyzed, and data output files.

