## Supplementary Material for "The differing prevalences of propionate and butyrate-producing bacteria in the human gut microbiota"

**Supplementary Text**

**Text S1 - Single-pathway determination for strains testing positive for multiple pathways:** 26 strains tested positive for both WWC and SP, including two model strains: Veillonella parvula and Propionibacterium acidipropionici. HMMER scores for these strains were consistently closer to model SP or WWC strains for the decarboxylase genes of the SP pathway than they were for the methylmalonyl-CoA decarboxylase of the WWC, with one exception: P. acidipropionici had higher scores for WWC. All strains were thus reclassified as SP strains except for P. acidipropionici, which was reclassified as a WWC strain. This method is consistent with model strains as V. parvula is a model SP strain and P. acidipropionici is a model WWC strain.

Three strains tested positive for both SP and Acr, all in the genus Veillonella. These strains were recategorized as SP because they had HMMER scores consistently within the 1st quartile for SP genes but ranged from below the 1st to the 2nd quartile for Acr genes. Additionally, V. parvula is a model strain for the SP pathway.

Several strains tested positive for multiple butyrate pathways: 12 strains for both Ace and 4-Ami, 19 for Ace and Lys, 18 for Ace and Glu, 4 for Lyc and 4-Ami, 5 for Lys and Glu, and 1 for 4-Ami and Glu. From a comparison of HMMER scores for genes for each pathway that these strains tested positive for, we recategorized all Ace and Glu positive strains as Glu, all Ace and 4-Ami strains as 4-Ami, all Lys and Glu strains as Lys, all Glu and 4-Ami strains as 4-Ami, and all 4-Ami and Lys strains as Lys. HMMER scores were similar between the Lys and Ace positive strains, so after the recategorization step, 11 strains remained categorized as both Lys and Ace.

**Text S2 - Metagenomic Datasets Included in this Study:** Five publicly available metagenomic datasets were included in this study for a total of 1,973 samples and 1,639 subjects (Asnicar et al., 2017, 2021; Bäckhed et al., 2015; Lloyd-Price et al., 2019; Schirmer et al., 2018) (PMID: 29311644, 31142855, 25974306, 28144631, 33432175). Subjects spanned across four countries, Italy, Great Britain, Sweden, and the United States, and ranged in age from newborn to 76 years old. 1,639 subjects had available data on gut health and were diagnosed as having healthy guts or Inflammatory Bowel Disease (IBD), which was further diagnosed as Crohn's Disease (CD) or Ulcerative Colitis (UC). N = 1,515 subjects had age data at the year or day level; n = 1,415 healthy individuals with available data on age. Based on previous studies showing significant gut microbiome differences among specific age groups (Asnicar et al., 2017; Ghosh et al., 2022), we further categorized subjects into the following age groups: 0 days to 2 weeks old (n = 90), 2 weeks to 6 months old (n = 102), 6 months to 3 years old (n = 98), 3 to 20 years old (n = 22), 20 to 60 years old (n = 995), and over 60 years old (n = 108).

### Supplementary Figures

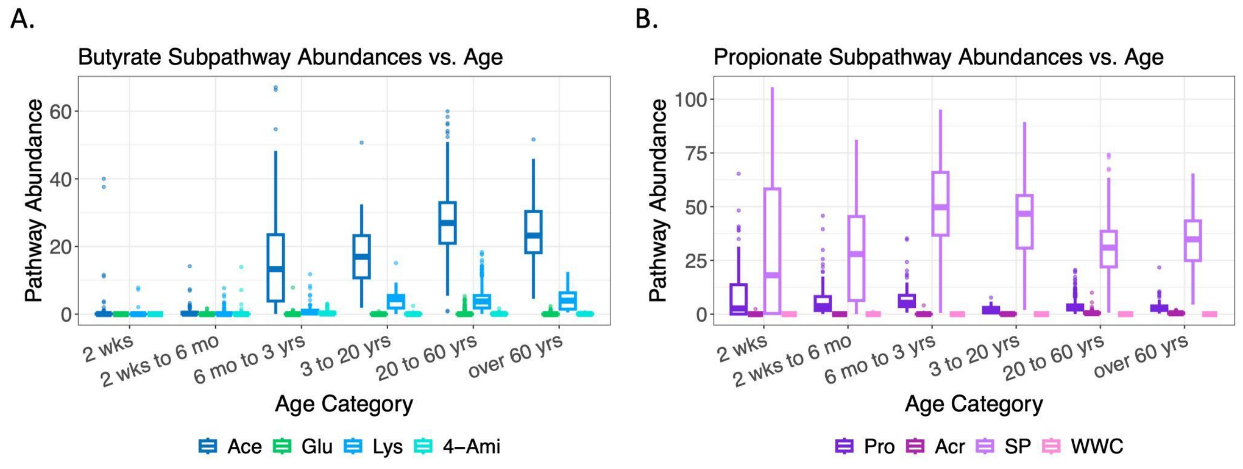


**Supplementary Figure S1. Variation in pathway abundance by age. A and B)** Detected abundance of each butyrate and propionate pathway by age category. Analyzed metagenomics datasets as described in **Fig. 3**.

**Supplementary Tables**

**Supplementary Table S1. Model strains used to determine required pathway genes and HMMER score cutoffs.**

| **pathway** | **model strain** |
|---|---|
| WWC | *Propionibacterium* spp. |
| SP | *Bacteroides fragilis* |
| SP | *Prevotella ruminicola* |
| SP | *Selenomonas ruminantium* |
| SP | *Veillonella parvula* |
| SP | *Selenomonas sputigena* |
| Acr | *Anaerotignum propionicum* |
| Acr | *Megasphaera elsdenii* |
| Pro | *Roseburia inulinivorans* |
| Pro | *Salmonella typhimurium* |
| Lys | *Clostridium sticklandii* |
| Lys | *Fusobacterium nucleatum* |
| Lys | *Porphyromonas gingivalis* |
| Lys | *Eubacterium ramulus* |
| Glu | *Acidaminococcus fermentans* |
| Glu | *Clostridium symbiosum* |
| Glu | *Peptoniphilus asaccharolyticus* |
| 4-Ami | *Anaerostipes rhamnosivorans* |
| Ace | *Anaerostipes caccae* |
| Ace | *Butyricicoccus pullicaecorum* |
| Ace | *Clostridium butyricum* |
| Ace | *Coprococcus catus* |
| Ace | *Eubacterium limosum* |
| Ace | *Eubacterium rectale* |
| Ace | *Megasphaera elsdenii* |

| | |
|---|---|
| Ace | *Roseburia hominis* |
| Ace | *Faecalibacterium prausnitzii* |
| Ace | *Anaerostipes hadrus* |
| Ace | *Anaerotruncus colihominis* |
| Ace | *Clostridium acetobutylicum* |
| Ace | *Coprococcus comes* |
| Ace | *Eubacterium ruminantium* |
| Ace | *Roseburia inulinivorans* |
| Ace | *Butyricicoccus pullicaecorum* |
| Ace | *Butyrivibrio fibrisolvens* |
| Ace | *Eubacterium ramulus* |
| Ace | *Roseburia faecis* |

**Supplementary Table S2. Probed gut species for which one of the eight propionate or butyrate producing pathways was detected.**

| **Strain** | **Pathway** |
|---|---|
| *Anaerofustis stercorihominis* | 4 Ami |
| *Anaerostipes caccae* DSM 14662 MAF 2 | 4 Ami |
| *Butyricicoccus pullicaecorum* | 4 Ami |
| *Clostridioides difficile* | 4 Ami |
| *Clostridium beijerinckii* | 4 Ami |
| *Clostridium difficile* | 4 Ami |
| *Clostridium populeti* | 4 Ami |
| *Clostridium sardiniense* | 4 Ami |
| *Clostridium symbiosum* | 4 Ami |
| *Anaerobutyricum hallii* | Ace |
| *Anaerococcus hydrogenalis* | Ace |
| *Anaerococcus murdochii* | Ace |
| *Anaerococcus obesiensis* | Ace |

| | |
|---|---|
| *Anaerococcus octavius* | Ace |
| *Anaerococcus prevotii* | Ace |
| *Anaerococcus senegalensis* | Ace |
| *Anaerococcus vaginalis* | Ace |
| *Anaerostipes butyraticus* | Ace |
| *Anaerostipes coli* | Ace |
| *Anaerostipes hadrus* | Ace |
| *Anaerotruncus colihominis* DSM 17241 MAF 2 | Ace |
| *Aneurinibacillus aneurinilyticus* | Ace |
| *Aneurinibacillus migulanus* | Ace |
| *Bacillus altitudinis* | Ace |
| *Bacillus amyloliquefaciens* | Ace |
| *Bacillus arsenicus* | Ace |
| *Bacillus atrophaeus* | Ace |
| *Bacillus badius* | Ace |
| *Bacillus beijingensis* | Ace |
| *Bacillus benzoevorans* | Ace |
| *Bacillus cereus* | Ace |
| *Bacillus endophyticus* | Ace |
| *Bacillus firmus* | Ace |
| *Bacillus flexus* | Ace |
| *Bacillus idriensis* | Ace |
| *Bacillus infantis* | Ace |
| *Bacillus licheniformis* | Ace |
| *Bacillus marisflavi* | Ace |
| *Bacillus massiliosenegalensis* | Ace |
| *Bacillus mojavensis* | Ace |
| *Bacillus mycoides* | Ace |
| *Bacillus niacini* | Ace |
| *Bacillus pumilus* | Ace |

| | |
|---|---|
| *Bacillus siralis* | Ace |
| *Bacillus sonorensis* | Ace |
| *Bacillus subtilis* | Ace |
| *Bacillus thuringiensis* | Ace |
| *Bacillus timonensis* | Ace |
| *Bacillus vallismortis* | Ace |
| *Bilophila wadsworthia* | Ace |
| *Brachymonas chironomi* | Ace |
| *Brachyspira pilosicoli* | Ace |
| *Brevibacterium halotolerans* | Ace |
| *Brevundimonas aurantiaca* | Ace |
| *Brevundimonas bullata* | Ace |
| *Brevundimonas diminuta* | Ace |
| *Brevundimonas terrae* | Ace |
| *Brevundimonas vesicularis* | Ace |
| *Butyricimonas faecihominis* | Ace |
| *Butyricimonas synergistica* | Ace |
| *Butyrivibrio crossotus* DSM 2876 MAF 2 | Ace |
| *Butyrivibrio fibrisolvens* | Ace |
| *Christensenella minuta* | Ace |
| *Chryseobacterium hominis* | Ace |
| *Cloacibacillus evryensis* | Ace |
| *Cloacibacterium normanense* | Ace |
| *Clostridium acetobutylicum* | Ace |
| *Clostridium aldenense* | Ace |
| *Clostridium baratii* | Ace |
| *Clostridium botulinum* | Ace |
| *Clostridium butyricum* | Ace |
| *Clostridium cadaveris* | Ace |
| *Clostridium celatum* | Ace |

| | |
|---|---|
| *Clostridium chauvoei* | Ace |
| *Clostridium cochlearium* | Ace |
| *Clostridium disporicum* | Ace |
| *Clostridium fallax* | Ace |
| *Clostridium felsineum* | Ace |
| *Clostridium indolis* | Ace |
| *Clostridium lituseburense* | Ace |
| *Clostridium methoxybenzovorans* | Ace |
| *Clostridium neonatale* | Ace |
| *Clostridium paraputrificum* | Ace |
| *Clostridium perfringens* | Ace |
| *Clostridium putrefaciens* | Ace |
| *Clostridium rectum* | Ace |
| *Clostridium saccharoperbutylacetonicum* | Ace |
| *Clostridium sartagoforme* | Ace |
| *Clostridium senegalense* | Ace |
| *Clostridium septicum* | Ace |
| *Clostridium sporogenes* | Ace |
| *Clostridium sticklandii* | Ace |
| *Clostridium tertium* | Ace |
| *Clostridium tyrobutyricum* | Ace |
| *Clostridium ventriculi* | Ace |
| *Clostridium vincentii* | Ace |
| *Coprococcus comes* ATCC 27758 MAF 2 | Ace |
| *Coprococcus eutactus* ATCC 27759 MAF 2 | Ace |
| *Coprococcus catus* | Ace |
| *Desemzia incerta* | Ace |
| *Desulfitobacterium frappieri* | Ace |
| *Eisenbergiella tayi* | Ace |
| *Empedobacter falsenii* | Ace |

| | |
|---|---|
| *Eubacterium ventriosum* ATCC 27560 MAF 2 | Ace |
| *Eubacterium barkeri* | Ace |
| *Eubacterium callanderi* | Ace |
| *Eubacterium cellulosolvens* | Ace |
| *Eubacterium contortum* | Ace |
| *Eubacterium desmolans* | Ace |
| *Eubacterium hadrum* | Ace |
| *Eubacterium hallii* | Ace |
| *Eubacterium limosum* | Ace |
| *Eubacterium moniliforme* | Ace |
| *Eubacterium oxidoreducens* | Ace |
| *Eubacterium ramulus* | Ace |
| *Eubacterium rectale* | Ace |
| *Eubacterium ruminantium* | Ace |
| *Eubacterium xylanophilum* | Ace |
| *Facklamia tabacinasalis* | Ace |
| *Faecalibacterium prausnitzii* | Ace |
| *Flavobacterium cheniae* | Ace |
| *Fusobacterium mortiferum* | Ace |
| *Fusobacterium necrogenes* | Ace |
| *Gemmiger formicilis* | Ace |
| *Geobacillus vulcani* | Ace |
| *Intestinimonas butyriciproducens* | Ace |
| *Lachnobacterium bovis* | Ace |
| *Megasphaera elsdenii* 14 14 MAF 2 | Ace |
| *Microbacterium schleiferi* | Ace |
| *Oscillibacter valericigenes* | Ace |
| *Papillibacter cinnamivorans* | Ace |
| *Parasporobacterium paucivorans* | Ace |
| *Peptococcus niger* | Ace |

| | |
|---|---|
| *Peptoniphilus duerdenii* | Ace |
| *Planococcus okeanokoites* | Ace |
| *Planococcus rifietoensis* | Ace |
| *Planomicrobium chinense* | Ace |
| *Pseudoramibacter alactolyticus* | Ace |
| *Rheinheimera perlucida* | Ace |
| *Rheinheimera texasensis* | Ace |
| *Riemerella columbina* | Ace |
| *Romboutsia lituseburensis* | Ace |
| *Roseburia intestinalis* L1 82 MAF 2 | Ace |
| *Roseburia faecis* | Ace |
| *Roseburia hominis* | Ace |
| *Roseburia inulinivorans* | Ace |
| *Rudanella lutea* | Ace |
| *Ruminococcus gauvreauii* DSM 19829 MAF 2 | Ace |
| *Sarcina ventriculi* | Ace |
| *Sporosarcina koreensis* | Ace |
| *Subdoligranulum variabile* DSM 15176 MAF 2 | Ace |
| *Thauera terpenica* | Ace |
| *Thermus scotoductus* | Ace |
| *Tissierella praeacuta* | Ace |
| *Variovorax boronicumulans* | Ace |
| *Virgibacillus proomii* | Ace |
| *Wautersiella falsenii* | Ace |
| *Acidaminococcus intestini* D21 MAF 2 | Glu |
| *Acidaminococcus fermentans* | Glu |
| *Clostridium sp M62 1 MAF 2* | Glu |
| *Filifactor alocis* | Glu |
| *Kallipyga massiliensis* | Glu |
| *Peptoniphilus asaccharolyticus* | Glu |

| | |
|---|---|
| *Peptoniphilus grossensis* | Glu |
| *Peptoniphilus harei* | Glu |
| *Peptoniphilus indolicus* | Glu |
| *Peptoniphilus lacrimalis* | Glu |
| *Peptoniphilus senegalensis* | Glu |
| *Peptoniphilus timonensis* | Glu |
| *Alistipes putredinis* DSM 17216 MAF 2 | Lys |
| *Butyricimonas virosa* DSM 23226 MAF 2 | Lys |
| *Butyricimonas faecihominis* | Lys |
| *Butyricimonas paravirosa* | Lys |
| *Butyricimonas synergistica* | Lys |
| *Cetobacterium somerae* | Lys |
| *Clostridium felsineum* | Lys |
| *Clostridium lituseburense* | Lys |
| *Clostridium putrefaciens* | Lys |
| *Clostridium senegalense* | Lys |
| *Clostridium sticklandii* | Lys |
| *Eubacterium ramulus* | Lys |
| *Flavonifractor plautii* | Lys |
| *Fusobacterium gonidiaformans* | Lys |
| *Fusobacterium necrophorum* | Lys |
| *Fusobacterium nucleatum* | Lys |
| *Fusobacterium periodonticum* | Lys |
| *Fusobacterium russii* | Lys |
| *Fusobacterium varium* | Lys |
| *Intestinimonas butyriciproducens* | Lys |
| *Lachnoanaerobaculum umeaense* | Lys |
| *Odoribacter laneus* | Lys |
| *Odoribacter splanchnicus* | Lys |
| *Porphyromonas asaccharolytica* | Lys |

| | |
|---|---|
| *Porphyromonas endodontalis* | Lys |
| *Porphyromonas gingivalis* | Lys |
| *Porphyromonas somerae* | Lys |
| *Porphyromonas uenonis* | Lys |
| *Romboutsia lituseburensis* | Lys |
| *Tissierella praeacuta* | Lys |
| *Tumebacillus permanentifrigoris* | Lys |
| *Anaerotignum propionicum* | Acr |
| *Clostridium lactatifermentans* | Acr |
| *Clostridium propionicum* | Acr |
| *Clostridium tyrobutyricum* | Acr |
| *Coprococcus catus* | Acr |
| *Fusobacterium necrophorum* | Acr |
| *Megasphaera elsdenii* 14 14 MAF 2 | Acr |
| *Megasphaera elsdenii* | Acr |
| *Papillibacter cinnamivorans* | Acr |
| *Peptoniphilus indolicus* | Acr |
| *Peptostreptococcus anaerobius* | Acr |
| *Blautia wexlerae* DSM 19850 MAF 2 | Pro |
| *Blautia luti* | Pro |
| *Citrobacter amalonaticus* | Pro |
| *Citrobacter braakii* | Pro |
| *Citrobacter farmeri* | Pro |
| *Citrobacter freundii* | Pro |
| *Citrobacter gillenii* | Pro |
| *Citrobacter koseri* | Pro |
| *Citrobacter murliniae* | Pro |
| *Citrobacter pasteurii* | Pro |
| *Citrobacter sedlakii* | Pro |
| *Citrobacter werkmanii* | Pro |

| | |
|---|---|
| *Citrobacter youngae* | Pro |
| *Clostridium disporicum* | Pro |
| *Clostridium fallax* | Pro |
| *Clostridium neonatale* | Pro |
| *Eisenbergiella tayi* | Pro |
| *Enterobacter massiliensis* | Pro |
| *Enterococcus avium* | Pro |
| *Escherichia albertii* | Pro |
| *Escherichia fergusonii* | Pro |
| *Eubacterium contortum* | Pro |
| *Fusobacterium nucleatum* | Pro |
| *Fusobacterium varium* | Pro |
| *Hafnia paralvei* | Pro |
| *Klebsiella oxytoca* | Pro |
| *Klebsiella pneumoniae* | Pro |
| *Lactobacillus brevis* | Pro |
| *Pediococcus acidilactici* | Pro |
| *Robinsoniella peoriensis* | Pro |
| *Roseburia inulinivorans strain* AF28 15 | Pro |
| *Roseburia inulinivorans* | Pro |
| *Ruminococcus faecis* | Pro |
| *Ruminococcus gnavus* | Pro |
| *Salmonella enterica* | Pro |
| *Salmonella typhimurium* | Pro |
| *Yersinia aleksiciae* | Pro |
| *Yersinia bercovieri* | Pro |
| *Yersinia enterocolitica* | Pro |
| *Yersinia frederiksenii* | Pro |
| *Yersinia kristensenii* | Pro |
| *Akkermansia muciniphila* | SP |

| | |
|---|---|
| *Alistipes indistinctus* YIT 12060 MAF 2 | SP |
| *Alistipes onderdonkii* DSM 19147 MAF 2 | SP |
| *Alistipes putredinis* DSM 17216 MAF 2 | SP |
| *Alistipes senegalensis* JC50 MAF 2 | SP |
| *Alistipes shahii* WAL 8301 MAF 2 | SP |
| *Alistipes finegoldii* | SP |
| *Alistipes obesi* | SP |
| *Alistipes timonensis* | SP |
| *Bacteroides caccae* ATCC 43185 MAF 2 | SP |
| *Bacteroides cellulosilyticus* DSM 14838 MAF 2 | SP |
| *Bacteroides coprophilus* DSM 18228 MAF 2 | SP |
| *Bacteroides dorei* 5 1 36 D4 MAF 2 | SP |
| *Bacteroides dorei* DSM 17855 MAF 2 | SP |
| *Bacteroides eggerthii* DSM 20697 MAF 2 | SP |
| *Bacteroides finegoldii* DSM 17565 MAF 2 | SP |
| *Bacteroides fragilis* 2 1 16 MAF 2 | SP |
| *Bacteroides fragilis* 3 1 12 MAF 2 | SP |
| *Bacteroides intestinalis* DSM 17393 MAF 2 | SP |
| *Bacteroides ovatus* ATCC 8483 MAF 2 | SP |
| *Bacteroides plebeius* DSM 17135 MAF 2 | SP |
| *Bacteroides rodentium* JCM 16496 MAF 2 | SP |
| *Bacteroides sp* 3 1 19 MAF 2 | SP |
| *Bacteroides sp* 9 1 42FAA MAF 2 | SP |
| *Bacteroides sp* D2 MAF 2 | SP |
| *Bacteroides stercoris* ATCC 43183 MAF 2 | SP |
| *Bacteroides thetaiotaomicron* 1 1 6 MAF 2 | SP |
| *Bacteroides thetaiotaomicron* VPI 5482 MAF 2 | SP |
| *Bacteroides uniformis* ATCC 8492 MAF 2 | SP |
| *Bacteroides xylanisolvens* SD CC 1b MAF 2 | SP |
| *Bacteroides acidifaciens* | SP |

| | |
|---|---|
| *Bacteroides barnesiae* | SP |
| *Bacteroides clarus* | SP |
| *Bacteroides coprocola* | SP |
| *Bacteroides faecichinchillae* | SP |
| *Bacteroides faecis* | SP |
| *Bacteroides fluxus* | SP |
| *Bacteroides fragilis* | SP |
| *Bacteroides gallinarum* | SP |
| *Bacteroides graminisolvens* | SP |
| *Bacteroides massiliensis* | SP |
| *Bacteroides nordii* | SP |
| *Bacteroides oleiciplenus* | SP |
| *Bacteroides pyogenes* | SP |
| *Bacteroides salanitronis* | SP |
| *Bacteroides salyersiae* | SP |
| *Bacteroides sartorii* | SP |
| *Bacteroides stercorirosoris* | SP |
| *Bacteroides thetaiotaomicron* | SP |
| *Bacteroides timonensis* | SP |
| *Bacteroides vulgatus* | SP |
| *Barnesiella intestinihominis* | SP |
| *Butyricimonas virosa* DSM 23226 MAF 2 | SP |
| *Butyricimonas faecihominis* | SP |
| *Butyricimonas paravirosa* | SP |
| *Butyricimonas synergistica* | SP |
| *Cloacibacillus evryensis* | SP |
| *Cloacibacillus porcorum* | SP |
| *Clostridium bifermentans* | SP |
| *Clostridium glycolicum* | SP |
| *Clostridium lituseburense* | SP |

| | |
|---|---|
| *Clostridium sordellii* | SP |
| *Clostridium sticklandii* | SP |
| *Coprobacter fastidiosus* | SP |
| *Coprobacter secundus* | SP |
| *Dialister invisus* | SP |
| *Dialister succinatiphilus* | SP |
| *Eubacterium tenue* | SP |
| *Flavonifractor plautii* | SP |
| *Intestinimonas butyriciproducens* | SP |
| *Megamonas funiformis* | SP |
| *Megamonas hypermegale* | SP |
| *Negativicoccus succinicivorans* | SP |
| *Odoribacter laneus* | SP |
| *Odoribacter splanchnicus* | SP |
| *Parabacteroides johnsonii* DSM 18315 MAF 2 | SP |
| *Parabacteroides merdae* ATCC 43184 MAF 2 | SP |
| *Parabacteroides sp D13 MAF 2* | SP |
| *Parabacteroides distasonis* | SP |
| *Parabacteroides faecis* | SP |
| *Parabacteroides goldsteinii* | SP |
| *Parabacteroides gordonii* | SP |
| *Paraprevotella clara* | SP |
| *Paraprevotella xylaniphila* | SP |
| *Phascolarctobacterium faecium* | SP |
| *Phascolarctobacterium succinatutens* | SP |
| *Phocaeicola vulgatus* | SP |
| *Porphyromonas asaccharolytica* | SP |
| *Porphyromonas endodontalis* | SP |
| *Porphyromonas gingivalis* | SP |
| *Porphyromonas uenonis* | SP |

| | |
|---|---|
| *Prevotella brevis* | SP |
| *Prevotella ruminicola* | SP |
| *Pyramidobacter piscolens* | SP |
| *Pyramidobacter piscolens* | SP |
| *Romboutsia lituseburensis* | SP |
| *Selenomonas noxia* | SP |
| *Selenomonas ruminantium* | SP |
| *Selenomonas sputigena* | SP |
| *Tannerella forsythia* | SP |
| *Tissierella praeacuta* | SP |
| *Veillonella sp* 3 1 44 MAF 2 | SP |
| *Veillonella sp* 6 1 27 MAF 2 | SP |
| *Veillonella atypica* | SP |
| *Veillonella dispar* | SP |
| *Veillonella parvula* | SP |
| *Veillonella ratti* | SP |
| *Veillonella rogosae* | SP |
| *Corynebacterium amycolatum* | WWC |
| *Corynebacterium durum* | WWC |
| *Corynebacterium glucuronolyticum* | WWC |
| *Corynebacterium kroppenstedtii* | WWC |
| *Corynebacterium ulcerans* | WWC |
| *Cutibacterium acnes* | WWC |
| *Propionibacterium acidipropionici* | WWC |
| *Propionibacterium acnes* | WWC |
| *Propionibacterium avidum* | WWC |
| *Propionibacterium freudenreichii* | WWC |
| *Propionibacterium jensenii* | WWC |
| *Propionimicrobium lymphophilum* | WWC |
